# Receptor interacting protein kinase-3 mediates both myopathy and cardiomyopathy in preclinical animal models of Duchenne muscular dystrophy

**DOI:** 10.1101/2022.01.06.475271

**Authors:** Maximilien Bencze, Baptiste Periou, Isabel Punzón, Inès Barthélémy, Valentina Taglietti, Cyrielle Hou, Louai Zaidan, Kaouthar Kefi, Stéphane Blot, Onnik Agbulut, Marianne Gervais, Geneviève Derumeaux, Laurent Tiret, François-Jérôme Authier, Fréderic Relaix

## Abstract

**Background:** Duchenne muscular dystrophy (DMD) is a progressive muscle degenerative disorder, culminating in a complete loss of ambulation, hypertrophic cardiomyopathy and a fatal cardiorespiratory failure.

Necroptosis is the form of necrosis that is dependent upon the receptor-interacting protein kinase (RIPK) 3; it is involved in several inflammatory and neurodegenerative conditions. We previously identified RIPK3 as a key player in the acute myonecrosis affecting the hindlimb muscles of the dystrophic mouse model, mdx. Whether necroptosis also mediates respiratory and heart disorders in DMD is currently unknown.

**Methods:** Evidence of activation of the necroptotic axis was examined in dystrophic tissues from Golden retriever muscular dystrophy (GRMD) dogs and R-DMDdel52 rats. A functional assessment of the involvement of necroptosis in dystrophic animals was performed on mdx mice that were genetically depleted for RIPK3. Dystrophic mice aged from 12 to 18 months were analyzed by histology and molecular biology to compare the phenotype of muscles from mdx*Ripk3*^+/+^ and mdx*Ripk3*^-/-^ mice. Heart function was also examined by echocardiography in 40-week-old mice.

**Results:** Quantification of *RIPK3* transcripts in sartorius and biceps femoris muscles from GRMD dogs positively correlated to myonecrosis levels (r=0.81; p=0.0076). *RIPK3* was also found elevated in the diaphragm (p=0<0.05). In the slow progressing heart phenotype of GRMD dogs, the phosphorylated form of RIPK1 at the Serine 161 site was dramatically increased in cardiomyocytes. A similar p-RIPK1 upregulation characterized the cardiomyocytes of R-DMDdel52 rats, associated with a marked overexpression *of Ripk1* (p=0.007) and *Ripk3* (p=0.008), indicating primed activation of the necroptotic pathway in the dystrophic heart. Mdx*Ripk3*^-/-^ mice displayed decreased compensatory hypertrophy of the heart (p=0.014), and echocardiography showed a 19% increase in the relative wall thickness (p<0.05) and 29% reduction in the left ventricle mass (p=0.0144). Besides, mdx*Ripk3*^-/-^ mice presented no evidence of a regenerative default or sarcopenia in skeletal muscles, moreover around 50% less affected by fibrosis (p<0.05).

**Conclusions:** Our data provide evidence of the activation of the necroptotic pathway in degenerative tissues from dystrophic animal models, including the diaphragm and the heart. The genetic inhibition of necroptosis in dystrophic mice improves both cardiac function and histological features of muscles, suggesting that prevention of necroptosis is susceptible to providing multiorgan beneficial effects for DMD.

## Introduction

Duchenne muscular dystrophy (DMD) is an X-linked genetic disorder, affecting 1/5000 male birth worldwide. Loss-of-function mutations in the dystrophin gene primarily lead to chronic myofibre necrosis (myonecrosis) and severe locomotor muscle disability starting in early childhood. Notably, the absence of dystrophin at the sarcolemma renders myofibre more susceptible to mechanical stress, which favours contraction-induced myonecrosis. Following injury, muscle repair requires myogenesis thanks to the contribution of PAX7-expressing resident muscle stem cells (MuSC)^1^. However, in disabling neuromuscular disorders involving chronic myofiber degeneration such as DMD, regeneration does not fully compensate for chronic myofibre demise. The loss of a sustained homeostatic balance between the two mechanisms leads to the increasing replacement of contractile myofibres by fibrotic or adipose tissue. Biopsies with interstitial fibrosis positively correlate with a poor outcome in DMD boys ^2^. Thus, identifying the molecular pathways involved in fibrosis formation is relevant for improving therapeutic perspectives. Dystrophin deficiency also challenges the survival of respiratory and cardiac muscle cells, leading to the premature death of DMD patients from cardiorespiratory weakness or failure ^3,4^.

Recently, we reported that necroptosis, a receptor-interacting protein kinase (RIPK)1-RIPK3-and mixed lineage domain kinase-like (MLKL)-dependent cell death pathway, is activated in the locomotor musculature of murine dystrophin-deficient muscles. Necroptosis inhibition dampens the peak of myonecrosis in the locomotor muscles of three-week-old mdx mice and improves muscle function in young adults ^5^. However, two recent studies reported a link between necroptosis inhibition and a deficit in early myogenesis and myofibre atrophy^6,7^. One of them reported that RIPK3 depletion in mdx MuSCs dramatically impairs muscle function. Repressing necroptosis might therefore lead to two putative conflictual outcomes: a decrease in myofibre degeneration, together with a default in muscle regeneration leading to muscle atrophy and sarcopenia. Consequently, the long-term effects of necroptosis inhibition in dystrophic skeletal and heart muscle remodelling need to be determined, before further envisaging the necroptotic pathway as a putative therapeutic target for DMD.

To this end, we examined the diaphragm and heart for evidence of necroptosis activation from Golden retriever muscular dystrophy (GRMD) dogs. The functional assessment of necroptosis in the cardiorespiratory system was achieved using mdx*Ripk3*^-/-^ mice. Compared to mdx*Ripk3*^+/+^, locomotor muscles from mdx*Ripk3*^-/-^ mice displayed a significant reduction of fibrosis. Despite its involvement in early myogenesis, RIPK3 deficiency did not generate sarcopenia. In the diaphragm, fibrosis reduction indicates that necroptosis not only participates in the pathology of limbs, but also of respiratory muscles. Importantly, we found that RIPK3 depletion improves cardiomyopathy by reducing heart hypertrophy and fibrosis deposition. Together, our data demonstrate that in dystrophic mdx mice, RIPK3 is dispensable for post-lesional myogenesis and suggests that necroptosis inhibition is beneficial on the hindlimb, respiratory, and heart muscles in dystrophinopathies. This work opens new avenues for further research exploring the improvement of patients’ health status by reducing necroptosis.

## Material and methods

### Animals

Mice and rats were bred in the Institut Mondor de Recherche biomédicale, Créteil, France in a pathogen-free facility with 12-h light and 12-h dark cycles in accordance with European Directive 2010/63/EU. Only male mice and rats were used for the experiments. Both wild-type (WT) and DMD rats and mdx mice were born from the same litter. The GRMD dogs included in this study were housed in the facilities of the neurobiology laboratory of the Veterinary School of Alfort. Experiments were approved by the Anses/EnvA/Upec Ethics Committee. All care and manipulations were performed by national and European legislation on animal experimentation. *Ripk3*^-/+^ mice were kindly provided by Genentech (San Francisco-CA). mdx*Ripk3*^-/-^ mice were generated by crossing *Ripk3^-/-^* with mdx mice. The resulting mice were backcrossed for at least four generations to obtain final mdx*Ripk3*^+/-^ breeders.

### Treadmill tests

The running capacity was evaluated on a treadmill (Tecmachine, Medical Development) at a +5% slope. Males of 70-to 76-week-old mice were initially acclimatized to the treadmill for 5 days (10 min/day and 12 m/min). Mice were challenged to an exercise of progressively increased intensity. Speed was increased every 90-to 120 seconds/min and the speed of the last completed running step before exhaustion was considered as the individual maximal aerobic speed.

### RNA extraction, PCR, and real-time PCR

Total skeletal muscle RNA was extracted from muscle and heart samples using TRIzol (Thermo Fisher Scientific) following the manufacturer’s instructions. SuperScript III Reverse Transcriptase from the Invitrogen kit converted RNA into cDNA using the Veriti 96-Well Fast Thermal Cycler (Applied Biosystems). Gene expression was quantified by real-time qPCR with the StepOnePlus real-time PCR system (Applied Biosystems) using SYBR Green detection tools (Applied Biosystems). In skeletal muscle tissue, the expression of each gene was normalized to *Tbp* gene expression. Results are reported as relative gene expression (2-DDCT). In the heart, from 500 ng of extracted RNA, the first-strand cDNA was then synthesized using a RevertAid First Strand cDNA Synthesis Kit (Thermo Fisher Scientific) with random hexamers according to the manufacturer’s instructions. Using the Light Cycler® 480 system (Roche Diagnostics), the reaction was carried out in duplicate for each sample in a 6-μl reaction volume containing 3 μl of SYBR Green Master Mix, 500 nM of the forward and reverse primers each and 3 μl of diluted (1:25) cDNA. The thermal profile for the SYBR Green qPCR was 95°C for 8 min, followed by 40 cycles at 95°C for 15 s, 60°C for 15 s and 72°C for 30 s. The mean gene expression was normalized to *Hprt1* and *Rpl32*. In dogs, RNA was extracted from frozen muscle slides using the NucleoSpin RNA extraction kit (Macherey Nagel). RNA was quantified by Nanodrop and the Maxima first strand cDNA master kit (Thermo Fisher) was used to convert 300ng of RNA into cDNA. Gene expression was quantified by real-time qPCR with the QuantStudio™ 3 Real-Time PCR System (Applied Biosystems) using SYBR Green detection kit (Thermo Fisher). The reaction was carried out in triplicates for each sample of diluted (1:10) cDNA. The expression of genes was normalized to *b-actin* gene expression. Results are reported as relative gene expression (2^-DDCT^).

Primers used: Mouse: forward *Ripk3* 5′-CGGGCACACCACAGAACAT-3′, *Ripk3* and reverse 5′-GTAGCACATCCCCAGCACCAC-3′, forward *Ripk1* 5′-AGAAGAAGGGAACTATTCGC-3′ and reverse *Ripk1* 5′-TTCTATGGCCTCCACGAT-3′, forward *Bnp* 5′-CAGCTCTTGAAGGACCAAGG-3′, reverse 5′-AGACCCAGGCAGAGTCAGAA-3′, forward *Myh7* 5′-AGGTGTGCTCTCCAGAATGG-3′, reverse 5′-CAGCGGCTTGATCTTGAAGT-3′, *Rpl32*-forward 5′-TGGTGAAGCCCAAGATCGTC-3′, reverse 5′-GGATCTGGCCCTTGAACCTT-3′, *Hprt1*-forward 5′-CAGGCCAGACTTTGTTGGAT-3′, reverse 5′-TTGCGCTCATCTTAGGCTTT-3′. Rat: forward *Ripk3* 5′-CGTACACGTAGTCCCACTCG-3′, reverse 5′-AGGGAGGTGAAGGCTATGGT-3′, forward *Ripk1* 5′-GGTCTCCCATGACCCCTTTG-3′, reverse 5′-GGTAGGTTGGTCTCAGGCAC-3′. Dog: Forward *Ripk1* 5′-GAATGAGTTCAGCCCTGCTC-3′, reverse 5′-CTCGCTCATAGTCGTGGTCA-3′, forward *Ripk3* 5′-CAGAGAGGCTCAAGGTCAGG-3′, reverse 5′-CGATGTCTGGGCCACTATCT-3′ *b-actin* forward 5′-CCATCTACGAGGGGTACGCCC-3′ reverse TGCTCGAAGTCC AGGGCGACGTA

### Histology

Fibrosis was labelled using Epredia Varistain Gemini ES Automated Slide Stainer. For immunofluorescence: rat antibody to CD68 (clone FA-11, 137001, 1/50), rabbit antibody to mouse pan-Laminin (Sigma, L9393, 1/1000), Phospho-RIPK1 (Ser161) antibody (1/100 Invitrogen 105640).

Biopsies from GRMD dogs were processed and analysed as previously described^8^. The dried sections were stained for 10 minutes in Hematein and 5 minutes in 1% Eosin, dehydrated in four consecutive baths of ethanol, one bath of xylene and mounted in Canada balsam. One entire section per biopsy was photographed using an AxioObserver Z1 linked to its ICC1 camera (Zeiss), and the MosaiX application of the software AxioVision (Zeiss). The image was then analysed using the software Visilog 6.4 (Noesis). A grid of 10 000 mm2 squares was superimposed onto the image of the entire section. At each intercept of the grid (i.e. every 100 mm) the histological aspect of the underlying tissue was manually captioned using predefined annotations. The percentage of each type of histological event was calculated, defined as the percentage of events not corresponding to normal shape fibres. Inflammatory infiltration in dog biopsies was quantified using CD4, CD8, and CD11b immunostainings. 7μm sections from each biopsy were fixed in cold acetone-methanol. After having blocked the endogenous peroxidase activity, the primary antibody (either a rat anti-canine CD4, Serotec H, 1/50, or a rat anti-canine CD8, Serotec H, 1/50, or a mouse anti-canine CD11b, Serotec H, 1/50) was used. The amount of inflammatory cells was normalized to the area of the section^8^.

Myonecrosis levels in mdx biopsies were determined by measuring the area corresponding to mouse IgG uptake in myofibres (and not in interstitial areas) and expressed as the percentage of the cross-sectional area. Muscle fibre minimal Feret diameter, corresponding to the minimum distance between the two parallel tangents of myofibres, was determined. Images analyses were performed with ImageJ by a specific self-developed macro that recognizes muscle fibres as previously described ^9^.

### Cardiac Phenotyping

Mice were trained to be grasped since transthoracic echocardiography (TTE) was performed in conscious, non-sedated mice to avoid any cardiac depressor effect of sedative or anaesthetic agents. Data acquisition was performed every month by a single operator (JT) as previously described ^10^. Images were acquired from a parasternal position at the level of the papillary muscles using a 13-MHz linear-array transducer with a digital ultrasound system (Vivid 7, GE Medical System, Horton, Norway). Left ventricular diameters and ejection fraction, anterior and posterior wall thicknesses were serially obtained from M-mode acquisition. Relative LV wall thickness (RWT) was defined as the sum of septal and posterior wall thickness over LV end-diastolic diameter, and LV mass was determined using the uncorrected cube assumption formula (LV mass=(IVSd+LVIDd+PWTd)^3^-LVIDd^3^). Peak systolic values of radial SR in the anterior and posterior wall were obtained using Tissue Doppler Imaging (TDI) as previously described ^10^. TDI loops were acquired from the same parasternal view at a mean frame rate of 514 fps and a depth of 1 cm. The Nyquist velocity limit was set at 12cm/s. Radial SR analysis was performed offline using the EchoPac Software (GE Medical Systems) by an observer (GD) blinded to the diet of the animals. The Peak systolic of radial SR was computed from a region of interest positioned in the mid anterior wall and was measured over an axial distance of 0.6 mm. The temporal smoothing filters were turned off for all measurements. Because slight respiratory variations exist, we averaged peak systolic of radial SR on 8 consecutive cardiac cycles.

## Results

Firstly, we addressed the relevance of necroptosis in dystrophin deficiency as a potential cell demise involved in locomotor and respiratory muscles and the heart. Because heart or diaphragm muscle biopsies cannot be performed in patients, we aimed at examining tissues in different animal models for DMD. We first investigated degenerating tissues from golden retriever muscular dystrophy (GRMD) dogs that represent a gold standard relevant animal model for DMD^11^. Compared to age-matched wild-type (WT) dogs, 6-month-old-GRMD dogs were characterized by typical high and variable blood creatine kinase levels (p=0.0167) (Figure 1a), assessing a significant ongoing, global myofibre demise. Histological features of myonecrosis in the sartorius and biceps femoris muscles confirmed the presence of recently injured area (Figure 1b). Because myofibre necroptosis comes with an increase of *Ripk3* transcripts in mdx TA muscles, and that RIPK3 upregulation sensitizes muscle cells to TNF-elicited necroptosis^12^, we examined whether there was a correlation between muscles presenting high histological necrotic extent and *RIPK3* expression in muscles of GRMD dogs. Sartorius muscles from 6-and 12-month-old GRMD dogs were together quantified by qPCR for *RIPK3* transcripts and quantified for evidence of necrotic events and inflammation in hematoxylin & eosin staining. Interestingly, *RIPK3* expression was highly correlated with the extent of myonecrosis (Pearson test, p=0.0076, r=0.8138, n=9 GRMD muscles) (Figure 1b), but not with the inflammatory area (p=0.1409) (Figure 1c), suggesting that *RIPK3* elevation is specific to necrotic demise, rather than post-necrotic events such as myeloid infiltrate. Similarly, *RIPK3* transcripts increased in the GRMD diaphragm compared to age-matched WT dogs, and *RIPK3* expression was also positively correlated with *RIPK3* expression in these muscles (p=0.0307) (Figure 1d-f). Thus, RIPK3-dependant cell death does not only involve ambulatory, but also the main respiratory musculature in the DMD pathogenic course. In the heart, whilst we observed no significant increase in the necrosome transcripts in the left ventricle of 6-month-old GRMD dogs (Figure 1g,h), we found a dramatic increase of immunoreactivity of GRMD dog’s cardiomyocytes to antibodies directed against phospho-RIPK1 (Ser161), indicating that RIPK1 is primed for either extrinsic apoptosis or canonical necroptosis (Figure 1i)^13^’^14^.

**Figure 1:**
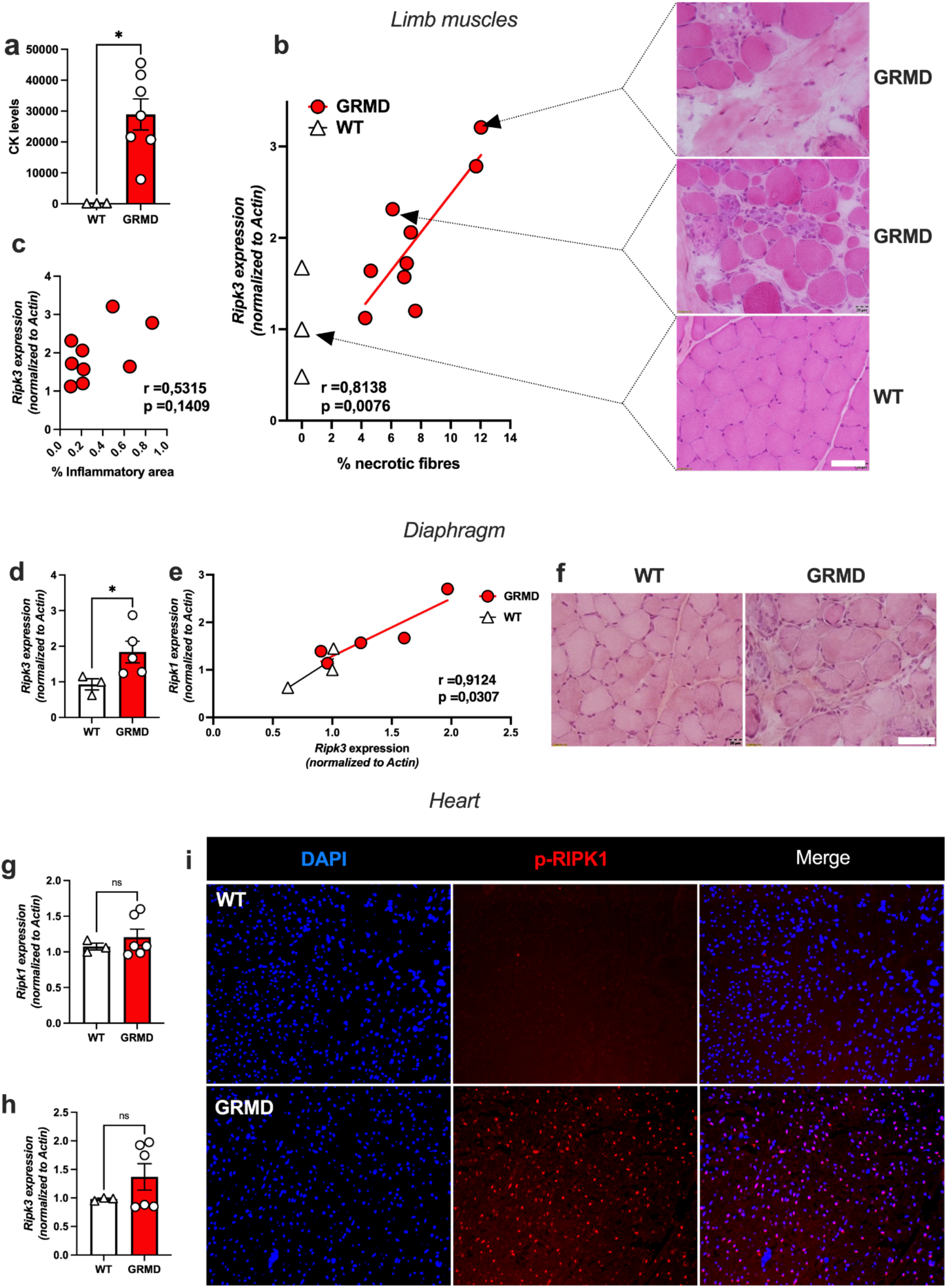
The RIPK1-RIPK3 axis is dysregulated in the skeletal muscles and the heart of dystrophic dogs. (a) Quantification of serum CK in six-month-old WT and GRMD dogs. (b) Pearson correlation coefficient (r) and p-value (p) between necrotic fibre percentage and *RIPK3* transcripts expression normalized to *ACTIN*. Representative H&E staining of the sartorius muscle corresponding to related biopsied muscle. Scale bar: 20 μm. (d) Pearson correlation coefficient and p-value between the percentage of inflammation and *RIPK3* transcripts expression. (d) Quantification of *RIPK3* expression in the diaphragms of 6-month-old WT and GRMD dogs. (e) Pearson correlation between *RIPK1* and *RIPK3* expression in the diaphragm. (f) H&E staining of the diaphragm muscle. (g) In the heart left ventricle of GRMD and WT dogs, quantification of *RIPK1* and (h) *RIPK3* transcripts normalized to *ACTIN*. (i) Representative image of a left ventricle heart from WT and GRMD dogs labelled with an antibody directed against the phosphorylated form of RIPK1 (S161). Data are given as means ± s.e.m. Ns: Not significant, *: p-value < 0.05.

Thus, to determine whether necroptosis has a functional role in progressive cardiomyopathy or in myonecrosis affecting the diaphragm of dystrophin-deficient mice, we generated mdx*Ripk3*^-/-^ mice by crossing mdx with *Ripk3-KO* mice, which compromise dystrophic mice for RIPK3-dependent necroptosis^15^. mdx*Ripk3*^+/+^ (hereafter named mdx*Ripk3*^+/+^ or mdx) and mdx*Ripk3*^-/-^ littermates were kept for ageing to reach the maximum mdx pathogenic state in striated muscles. We first examined whether the long-term genetic inhibition of necroptosis would promote sarcopenia by repressing MuSCs density and the regenerative capacity of skeletal muscle. Indeed, a recent article indicated that necroptosis inhibition generates a MuSCs depletion by inducing the death of PAX7-positive cells, with a subsequent deficit in myogenesis associated with a dramatically reduced muscle strength in adult mdx mice ^6^. In biceps from 18-month-old mdx*Ripk3*^-/-^ mice, *Ripk1* transcripts were downregulated compared to *Ripk3*^+/+^ littermates (Figure 2a), indicating that canonical necroptosis is repressed in mdx*Ripk3*^-/-^ mice. To determine whether the defect of necroptosis signalling compromises muscle function, mdx and mdx*Ripk3*^-/-^ littermates underwent forced treadmill exercise with progressive increase of speed, until exhaustion. Control mdx mice were characterized by an important variability in running performance, expressed in the running distance (Figure 2b) or the maximum aerobic speed before exhaustion (Figure 2c). mdx*Ripk3*^-/-^ mice showed more homogenous capacities (coefficient of variation: mdx: 20,99% *versus mdxRipk3*^-/-^ 12.66%). On average, mdx*Ripk3*^-/-^ ran ~ 10% longer than mdx littermates, despite no statistical difference likely due to the variability of the control mdx phenotype (p=0.5530). The putative atrophy of *Ripk3-/-* dystrophic muscles was examined using muscle mass normalized to the individual size of the animals, assessed by the tibial length (TL). TA, EDL and biceps muscle mass remained unchanged in aged mdx*Ripk3*^-/-^ compared to mdx*Ripk3*^+/+^ mice (Figure 2d). Although young mdx muscles show extensive necrotic events leading to the accumulation of fibrotic tissue, muscles from old mice are typically spared. In the rare post-necrotic areas in TAs from old mdx and mdx*Ripk3*^-/-^ mice, we found an equal density of Ki67-positive cells (Figure 2e,f), suggesting that necroptosis inhibition does not affect MuSC proliferation in mdx muscle at this age. The mean of pooled minimum Feret of TA myofibres (Figure 2g), as well as the relative distribution of myofibre size (Figure 2h), were strictly unchanged. Interestingly, we found that RIPK3 depletion reduced fibrosis in the TA (p=0.0252) (Figure 2i), suggesting a cumulative cytoprotective effect regarding ancient degenerative events. To further examine any possible evidence of regenerative default in mdx*Ripk3*^-/-^ muscles, we analysed biceps muscles for *Pax7* (Figure 2j) and *myogenin* (Figure 2k) expression but observed no difference. Of note, we found a significant downregulation of *myostatin* expression (p=0.0082), a biomarker of neuromuscular disorders while *follistatin* was not significantly repressed (p=0.4452) (Figure 2l,m).

**Figure 2:**
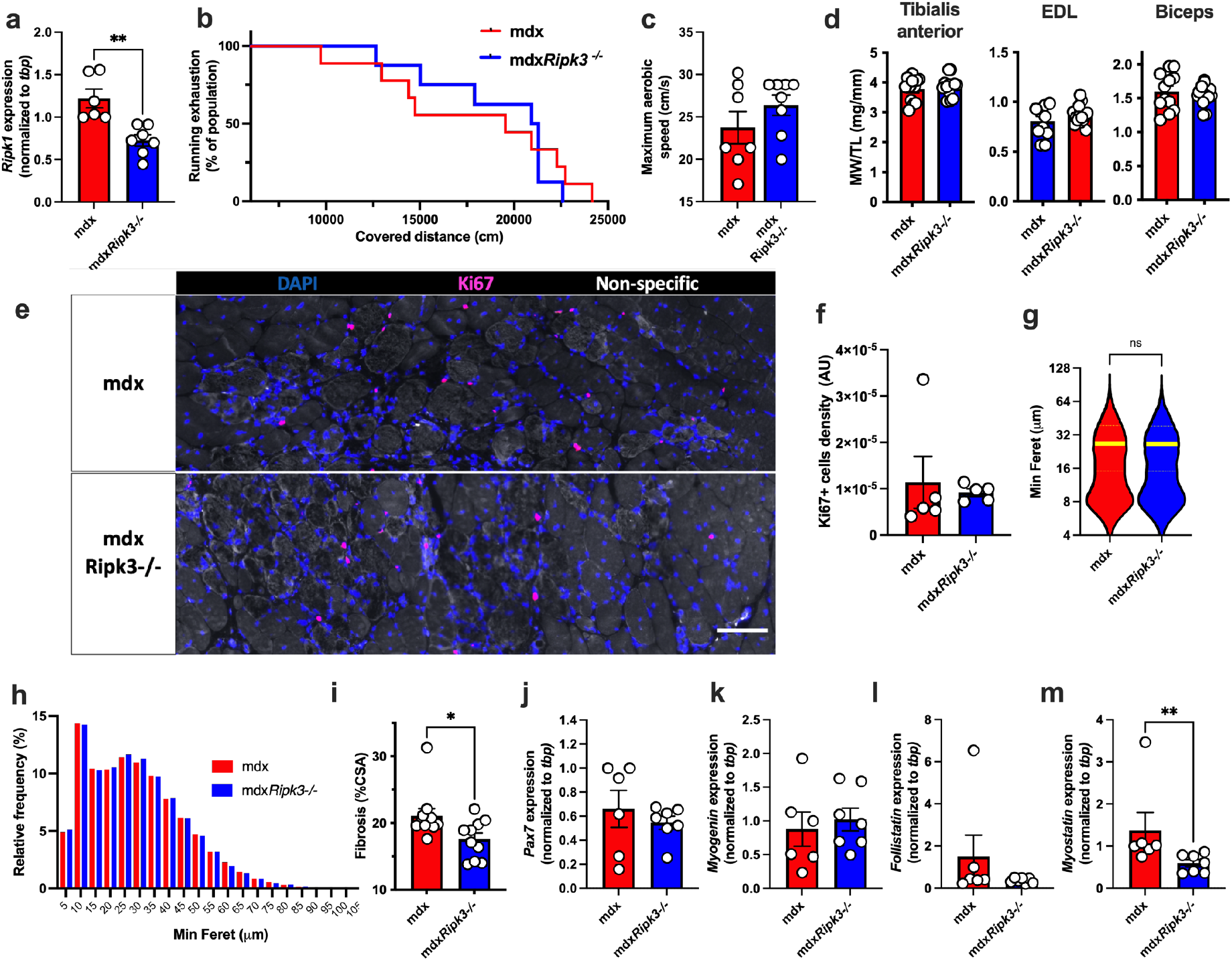
The genetic ablation of RIPK3 does not promote sarcopenia in 18-month-old mdx mice. Quantification of *Ripk1* transcripts in the eighteen-month-old biceps muscles (a) Seventeen-month-old *mdxRipk3+/+* (i.e. mdx) and *mdxRipk3-/-* male mice were submitted to forced treadmill running with a progressive increase of speed. (b) Endurance curve of running mice against covered distance expressed in cm. (c) Maximum aerobic speed before mice exhaustion. Data are given as means ± s.e.m. *mdxRipk3+/+*, n=7, *mdxRipk3-/-*, n=8. Shapiro-Wilk test followed by a Mann-Whitney test. (d) Tibialis anterior (TA), EDL and biceps muscles were harvested, and muscle weight (MW) was determined (and normalized to tibial length (MW/TL). (e) TA immunolabeling using antibodies directed against Ki67 (red). (f) Quantification of the density of Ki67-positive cells. (g) Violin plot of the pooled minimum Feret of TA myofibres (mdx: n=33,142 myofibres out of 7 distinct muscles; *mdxRipk3-/-*, n=26,469 myofibres out of 9 distinct muscles, Mann-Whitney test p=0.1526). (h) Relative frequencies of minimum Feret from TA myofibres are expressed in percentage. (i) Quantification of collagen deposition using Sirius red dye. The normal distribution of data was tested using the Kolmogorov-Smirnov test (passing normality test with α=0.05), leading to either t-tests or Mann-Whitney tests. (j) Quantification of the transcript levels of *Pax7*, (k), *myogenin*, (l) *follistatin* and (m) *myostatin* in the biceps, by quantitative PCR. Data are given as means ± s.e.m. Ns: Not significant, *: p-value < 0.05, **: p-value < 0.01. Scale bar:100 μm

We then examined whether the pathogenic features of the mdx diaphragm are improved by RIPK3 depletion (Figure 3). In histology, the localization of blood proteins such as albumin or immunoglobulins within sarcoplasm is a reliable marker for the necrotic fate of myofibre^16,17^. Muscles from eighteen-month-old mice showed IgG immunoreactivity almost only localized at the extracellular compartment (Figure 3a), and rarely within myofibres, indicating no/little ongoing myonecrosis. No difference was found in both genotypes (unpaired t-test, p=0.1914). The density of myofibres with leaky sarcolemma was extremely low in both genotypes (2.3 fibres/mm^2^ in mdx versus 2.6 fibres/mm^2^ in *mdxRipk3^-/-^)* (Figure 3b,c). This was neither associated with high-density cell pockets (Figure 3d) nor with CD68-positive cell infiltrates (Figure 3e). Furthermore, only a low percentage of centrally nucleated fibres was found in the mdx diaphragm (5-10% of myofibres) (Figure 3f). These observations suggested that the diaphragm from 18-month-old mdx mice stand at relative homeostasis with little/no ongoing myonecrosis, regardless of RIPK3 expression. Examining the profile of myofibre size, we observed a moderate decrease of myofibre size in mdx*Ripk3*^-/-^ diaphragm compared to mdx littermates (16.6 in mdx versus 16.1 in mdx*Ripk3*^-/-^, p<0.0001, Mann Whitney test, n>16,000 myofibres in each group, included in six mdx muscles, and eight mdx*Ripk3*^-/-^ muscles). Dystrophin-deficient muscles are characterized by an increased heterogeneity in myofibre size, illustrated by an increase of the variance coefficient^18^. RIPK3-deficiency decreased the mean-variance coefficient of myofibres in mdx mice (Figure 3i, unpaired t-test p=0.0431). Fibrosis deposition in respiratory muscles, such as the diaphragm, is a key pathogenic event in DMD pathogenesis and correlates with a bad prognosis in patients. To determine whether RIPK3 also mediates fibrosis deposition in the diaphragm, muscle sections were stained with Sirius red (Figure 3k), and we observed less collagen deposition in mdx*Ripk3*^-/-^diaphragms compared to mdx littermates (Figure 3k, unpaired t-test, p=0.0414). Together, our data demonstrate that RIPK3 participates in diaphragm pathogenesis in mdx mice by promoting interstitial fibrosis.

**Figure 3:**
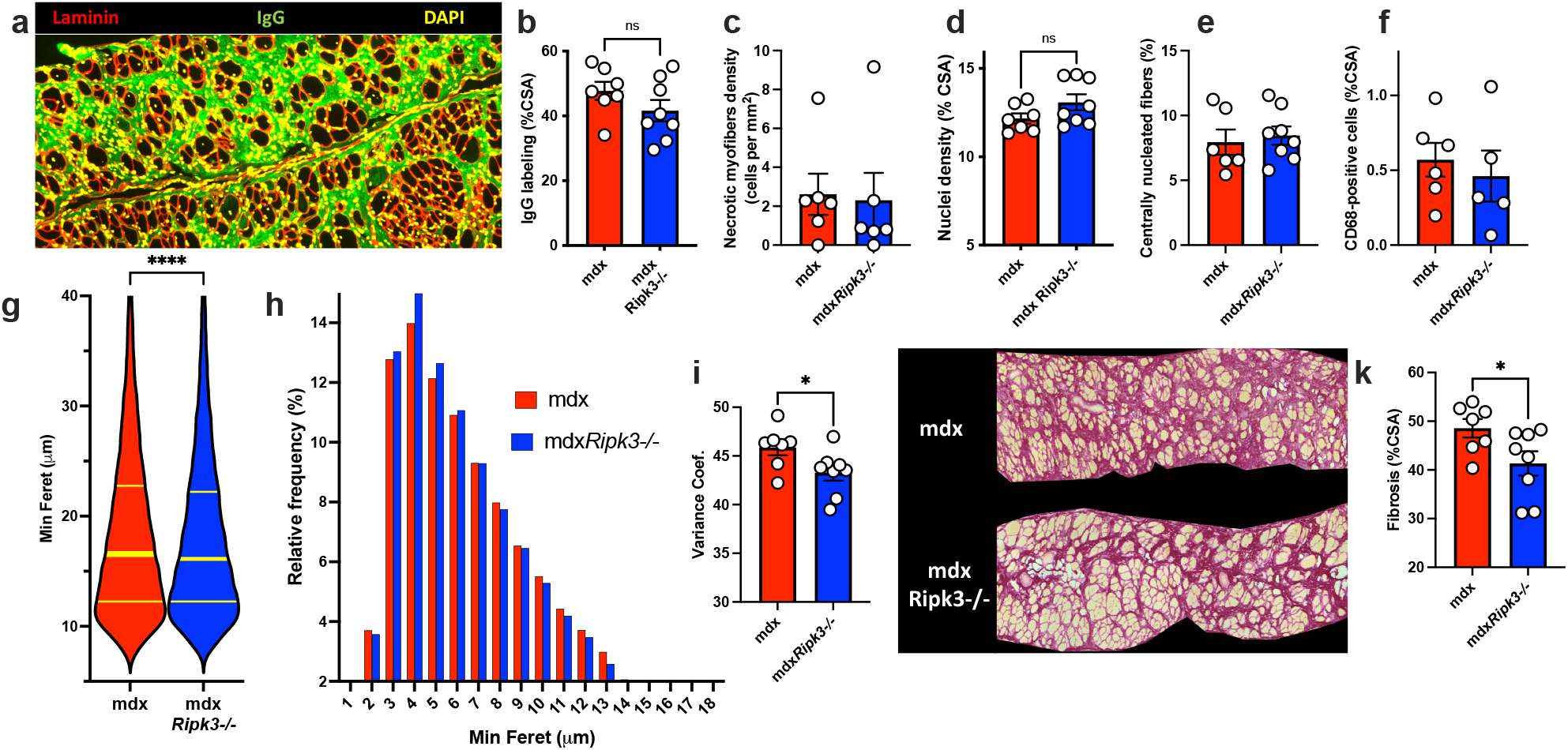
Necroptosis participates in the diaphragm fibrosis in mdx mice. (a) Representative image of IgG uptake in diaphragms of 18-month-old mdx (mdx*Ripk3*^+/+^) and *mdxRipk3~^/^~* mice. (b) Quantification of IgG uptake labelling expressed in percentage of crosssectional area (%CSA). (c) Quantification of myonecrosis extent. Myonecrosis is defined as the area occupied by leaky myofibres (i.e immunoreactive for mouse IgG uptake). Quantification of nuclear density expressed as a percentage of CSA (d), CD68-positive macrophage infiltrates expressed in percentage of CSA (e), and centrally nucleated myofibres expressed as a percentage of myofibres (f). (g) Violin plot of the minimum Ferets of diaphragm myofibres. (mdx, n=16873 including 7 distinct muscles, mean:18,61 ± 0.06555; *mdxRipk3-/-*, n=31666 including 8 distinct muscles, mean:18,21 ± 0.04554, two-tailed Mann-Whitney test p < 0,0001). (h) Relative frequency of minimum Ferets from diaphragm myofibres. Data are expressed as a percentage. (i) Quantification of the variance coefficient of the minimum Feret of myofibres, (j) representative picture of Sirius red staining (k) Quantification of diaphragm fibrosis. The normal distribution of data was tested using the Kolmogorov-Smirnov test (passing normality test with α=0.05), leading to either t-tests or Mann-Whitney tests. Data are given as means ± s.e.m. Ns: Not significant, *: p-value < 0.05, ****: p-value < 0.0001.

We next addressed the role of necroptosis in the development of cardiomyopathy upon dystrophin deficiency. Importantly, end-stage cardiomyopathy is a major concern in the survival of boys and is associated with hallmarks of cardiomyoblast necrosis and fibrosis deposition. In GRMD dogs, RIPK1 was found phosphorylated at the Ser161 site in the heart (Figure 1i), indicating an early step of activation of RIPK1-dependant apoptosis or necroptosis^14^. Similar p-RIPK1 immunoreactivity to GRMD dogs was observed in the hearts of R-DMDdel52 rats (Figure 4a) that develop an early cardiomyopathy^19^. Dystrophin-deficient rats also had a significant increase of *Ripk1* (p=0.0070, t-test) and *Ripk3* (p=0.0080, t-test) transcripts in heart tissue compared to WT rats (Figure 4b,c), suggesting advanced necroptosis activation in the heart from R-DMDdel52 rats. This contrasted with dogs that displayed no significant elevation of *RIPK1* and *RIPK3* transcription (Figure 1g,h), in agreement with rare fresh necrotic events observed in dystrophic hearts of GRMD dogs. To determine the role of necroptosis in the progressive cardiomyopathy of dystrophin-deficient hearts, we analysed the heart phenotype of 18-month-old mdx*Ripk3*^-/-^ mice (Figure 4 d-k). RIPK3 depletion decreased cardiac compensatory hypertrophy in mdx mice (Figure 4d,e). At the investigated age, we found no significant levels of ongoing degeneration in the heart, displaying little CD68-positive cell infiltration whatever the genotype (Figure 3f), which suggested an accumulation of past degenerative events with little evolution over time at this age. Interstitial collagen deposition was reduced by almost 50% in the right ventricle of mdx*Ripk3*^-/-^ mice compared to their mdx control littermates (Figure 4h,g). Furthermore, the hearts of mdx*Ripk3*^-/-^ mice showed significantly reduced levels of *Bnp* and *Myf7* transcripts, two sensitive markers of cardiac hypertrophy (Figure 4i,j).

**Figure 4:**
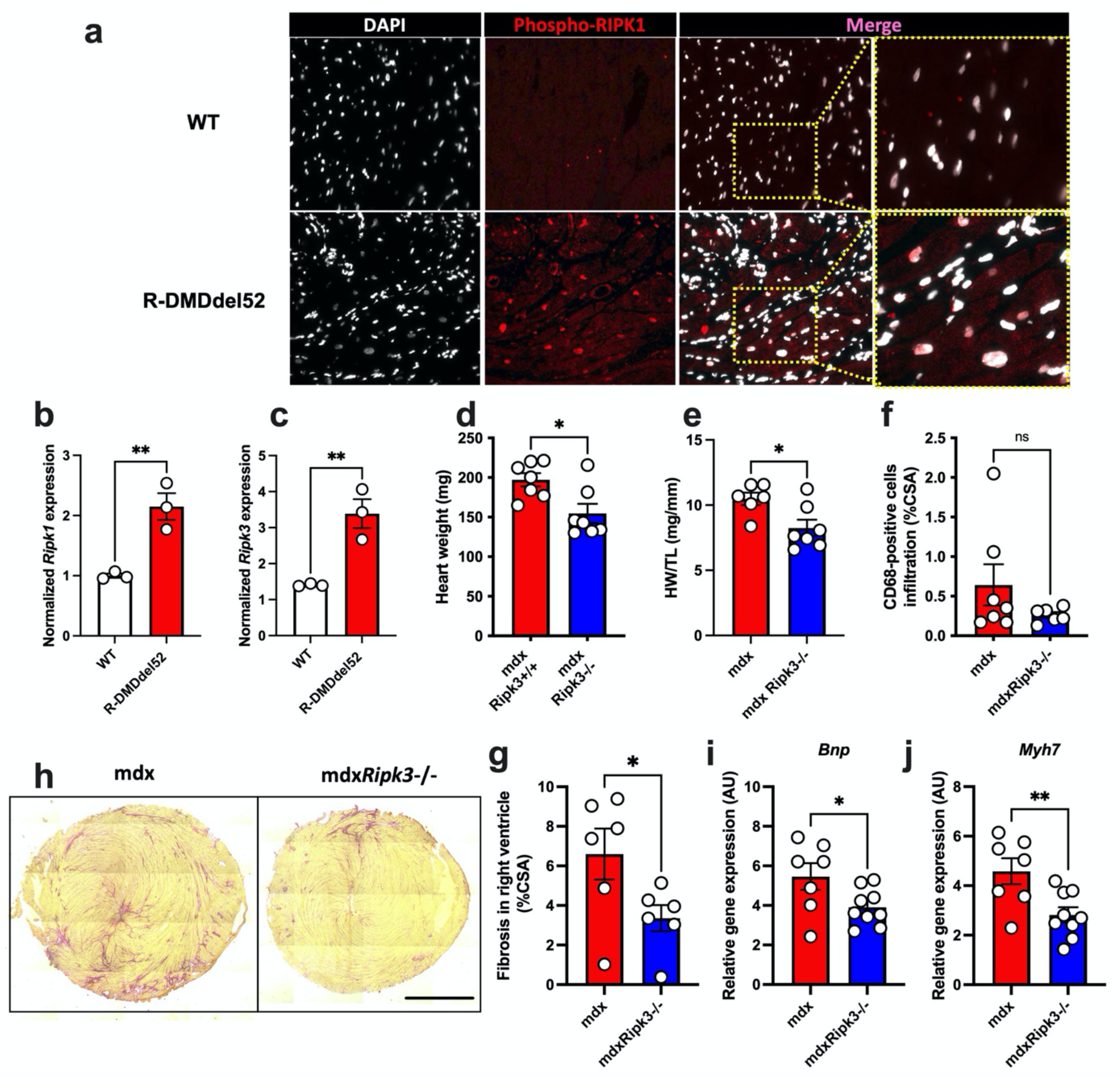
RIPK3 mediates the compensatory hypertrophy of the dystrophin-deficient heart. (a) Representative image of IgG uptake in diaphragms of 18-month-old mdx (mdx*Ripk3*^+/+^) and *mdxRipk3~^/^~* mice. (b) Quantification of IgG uptake labelling expressed in percentage of crosssectional area (%CSA). (c) Quantification of myonecrosis extent. Myonecrosis is defined as the area occupied by leaky myofibres (i.e immunoreactive for mouse IgG uptake). Quantification of nuclear density expressed as a percentage of CSA (d), CD68-positive macrophage infiltrates expressed in percentage of CSA (e), and centrally nucleated myofibres expressed as a percentage of myofibres (f). (a) Representative pictures of heart labelling using an antibody directed against the phosphorylated form of RIPK1 at S161 from wild-type (WT) and dystrophin-deficient R-DMDdel52 rats. Quantification of the rat *Ripk1* (d) and *Ripk3* (c) transcripts in hearts from WT and R-DMDdel52 rats. (d) Heart weights of 18-month-old mdx mice. (e) Heart weights normalized to the tibial length (TL). (f) Quantification of CD68-positive macrophage infiltrates. (g) Quantification of the density of CD31-positive microvessels, (h) representative picture of Sirius red staining of mdx and *mdxRipk3^-/-^*. Scale bar: 1700 μm. (i) Quantification of RV fibrosis. Hearts from 18-month-old mdx*Ripk3*^+l+^ and -/- were analyzed for *Bnp* (j), and *Myh7* (k) mRNA levels by quantitative PCR. The normal distribution of data was tested using the Kolmogorov-Smirnov test (passing normality test with α=0.05), leading to either t-tests or Mann-Whitney tests. Data are given as means ± s.e.m. Ns: Not significant, *: p-value < 0.05, **: p-value < 0.01.

We thus examined whether the improvement of the deleterious phenotype of cardiomyopathy conferred by RIPK3 depletion was validated at the 2D and functional levels and quantified routine echocardiographic parameters using the M(otion)-mode (Figure 5). Compared to wild-type mice, aged mdx mice are generally characterized by eccentric hypertrophy characterized by a transiently increased LV mass with increased left ventricle (LV) end-diastolic and end-systolic diameter. This is followed by a decompensated dilated cardiomyopathy associated with a reduction in LV fractional shortening and a decreased LV mass^20^. We confirmed that our 18-month-old mdx mice suffered from eccentric hypertrophy with an increased LV mass, associated with a reduced LV internal diameter in diastole (LVIDd) and systole (LVIDs) (Figure 5a-4c and Supplementary Table 1). Of note, compared to mdx mice, mdx*Ripk3*^-/-^ littermates displayed a 30% reduction of the LV mass linked to a decrease in LVIDd that contributed to the restoration of the LV relative wall thickness value (Figure 5d), and unchanged fractional shortening of the LV, strain rate of cardiomyocytes from the anterior wall (Figures 5e, f) and heart rate (Supplementary Table 1). Altogether, these data suggested that in the mdx model, RIPK3-dependent necroptosis participates in the long-term development of cardiac hypertrophy and dysfunction and that in these conditions, RIPK3 deficiency has overall cytoprotective effects against dystrophic-associated cardiac deleterious phenotype in the mouse.

**Figure 5:**
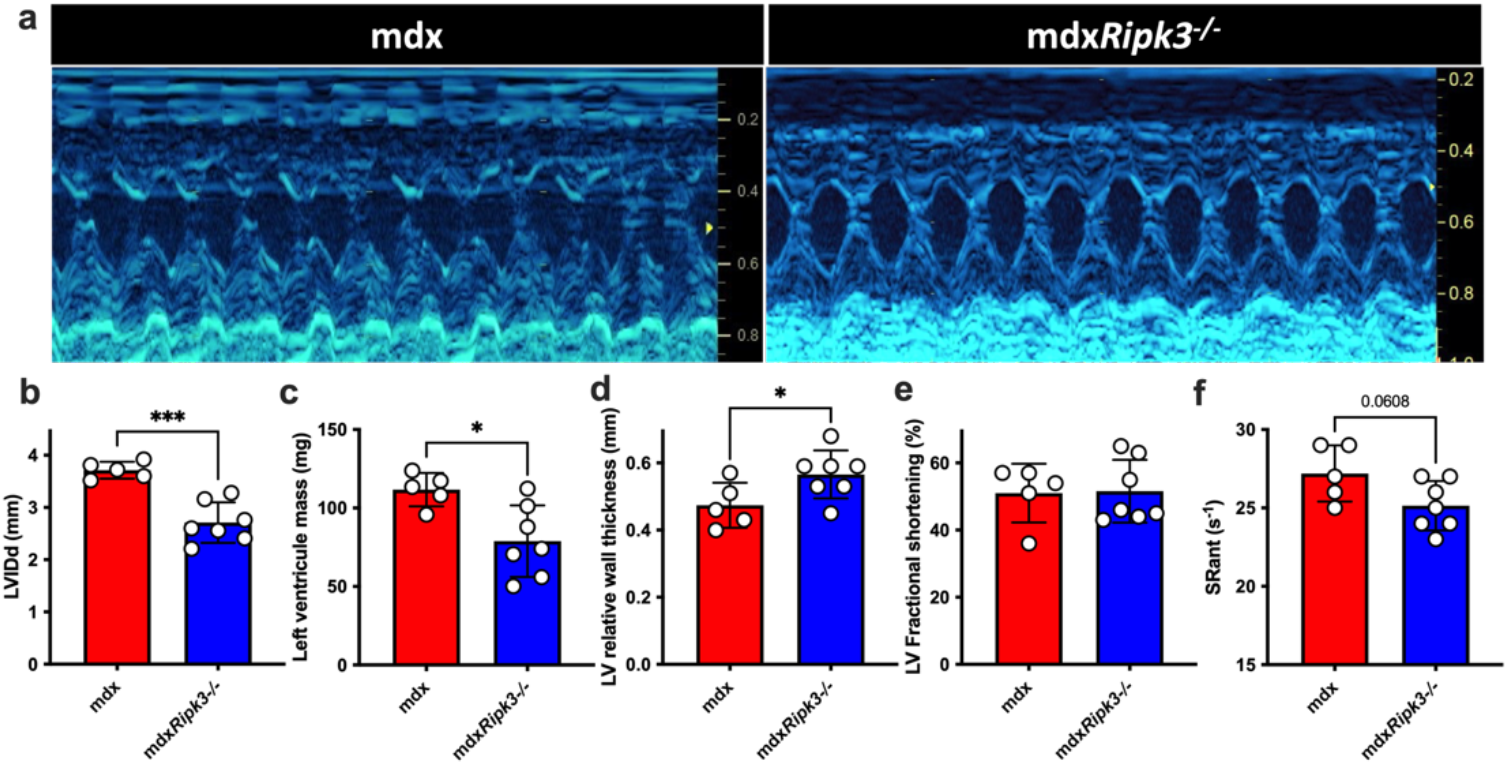
The genetic ablation of RIPK3 reduces cardiomyopathy in mdx mice. (a) Representative M-mode echocardiogram traces in mdx and mdx*Ripk3*^-/-^ 40-week-old mice showing a reduced left ventricle (LV) dilation in mdx*Ripk3*^-/-^ mice. (b) Quantification of LVIDd, (c) LV mass, (d) LV relative wall thickness, (e), LV fractional shortening, and (f) strain rate (SR) of the anterior LV wall. The normal distribution of data was tested using the Kolmogorov-Smirnov test (passing normality test with α=0.05), leading to t-tests. Data are given as means ± s.e.m. *: p-value < 0.05, ***: p-value < 0.001.

## Discussion

Necroptosis is now considered a new therapeutic target for several degenerating disorders, including cardiovascular neurodegenerative, and autoimmune diseases^21^. Dystrophin absence and DUX4 overexpression can lead to myofibre necroptosis *in vivo^12^*. Translating these discoveries into the clinic for neuromuscular disorders requires an understanding of 1-the long-term consequences of necroptosis inhibition in dystrophic conditions and 2-the role of necroptosis in the DMD cardiorespiratory phenotype.

While locomotor tissues can be harvested in DMD patients for diagnostic purposes and could be used for research purposes, respiratory or heart biopsies were not available. We then used dystrophic dogs and rats which are highly relevant animal models for DMD and reproduce the main aspects of DMD pathogenesis^11^. RIPK3 overexpression is a biomarker for myofibre necroptosis in mdx mice, and sensitizes myoblasts to TNF-elicited necroptosis^5^. Using sartorius and femoral biceps, diaphragm and cardiac tissues from 6-month-old GRMD dogs, we first assessed different levels of activation of the early steps of the RIPK1-RIPK3 canonical necroptotic pathway. RIPK3 upregulation in limb muscles, and diaphragm, and its correlation with the myonecrosis extent (Figure 1b), suggest a role for RIPK3 in the myonecrosis affecting GRMD dogs, especially in cells expressing p-RIPK1, which prime cells to programmed necrosis^14^. RIPK3 typically recruits MLKL during final necroptosis induction, and p-MLKL causes membrane permeability and necrotic morphology. Interestingly, dogs and other species belonging to the *Carnivora* order express crucial proteins required for necroptosis activation such as RIPK1, RIPK3, DNA-dependant activator of interferon regulatory factors (DAI) and TIR-domain-containing adaptor-inducing IFNβ (TRIF), however, they do not express MLK^23^. The molecular mechanisms of cell death execution in RIPK3-dependant but MLKL-independent necrosis, involvement remains an open question.

Notably, necrotic demise affecting the diaphragm and heart are key events leading to the short life expectancy of DMD boys. Herein, we provide evidence that beyond its role in the degeneration of locomotor muscles, necroptosis also takes part in the respiratory and cardiac phenotypes in the mdx model of DMD, notably by promoting tissue fibrosis. Accumulation of extracellular matrix deposition following myonecrosis is a central pathogenic process responsible for the loss of function of dystrophic muscles, by progressively replacing functional muscle tissue in both the myocardium and skeletal muscles^2^. Therefore, understanding the molecular pathways directly or indirectly leading to myofibre or cardiomyocyte death or fibrosis is highly relevant.

In dystrophin-deficient mice, investigating the phenotypes of the diaphragm and heart requires the generation of dystrophic and necroptosis-incompetent mice that would reach sufficient age to progressively develop a significant phenotype^24,25^. Compared to young dystrophic TA, the diaphragm from 18-month-old mdx mice showed IgG immunoreactivity mainly trapped in the large fibrotic area (Figure 3a) and barely detected within the myofibre compartment (Figure 3c), suggesting little/no ongoing myonecrosis. The observed low percentage of central nucleation together with low cell infiltration and high fibrosis indicates that this phenotype is the outcome of accumulating necrotic and regenerative events over time. Fibrosis decrease was observed in the diaphragm and was associated with a decrease in the mean myofibres size and size variability (Figure 3g-k). These results are consistent with a previous report assessing significant myofibre cytoprotection in the diaphragm^26^. We also found a significant reduction of fibrosis in the TA of old mdx*Ripk3*^-/-^ (Figure 2m) compared to mdx littermates, suggesting that the effects of necroptosis inhibition on muscle histology benefit both locomotor and respiratory muscles.

Two recent studies reported that necroptosis is involved in post-lesional myogenesis^6,27^. However, the underlying mechanisms of action of necroptosis on regeneration seem unclear: Sreenivasan and colleagues observed exclusive expression of RIPK3 in mdx MuSCs and not in myofibres. MuSCs depleted for RIPK3 led to a dramatic loss of PAX7-expressing cells and severely impaired muscle function^6^. On another hand, Zhou and colleagues found that myofibres express RIPK3 and MLKL and are susceptible to necroptotic demise. They demonstrated that myofibre necroptosis indirectly modulates muscle regeneration by releasing Tenascine-C, which is required for MuSCs proliferation. An effect of myofibre necroptosis on *Pax7* expression and myofibre size was observed *in vivo* at 7 and 15 days post-injury^27^. As the full effects on regeneration are generally observed at a later stage, after 4 weeks post-injury, it is unclear in this study whether the depletion of necroptosis irreversibly stops myogenesis or only delays it.

Herein, we found no impairment of running performance in dystrophic *Ripk3*^-/-^ mice, and no/little effect on muscle weight, while confirming the downregulation of *Ripk1* in muscle extracts. The weight of limb muscles was unchanged, as myofibre size (Figure 2a-h). *Pax7* and *myogenin* expression were not reduced in mdxRipk3-/-mice, suggesting there is no depletion of MuSCs. Together, our results demonstrate that in mdx mice, a long-term necroptosis inhibition does not lead to sarcopenia of dystrophic muscle, but rather has beneficial effects since it dampens fibrosis. The lack of observed sarcopenia in our mdx*Ripk3*^-/-^ dystrophic mice could for instance be explained by a delay of myogenesis when necroptosis is prevented. Indeed, slowing down the kinetics of differentiation does not necessarily impair regeneration^28^. It is also possible that any effect, positive or negative of RIPK3-deficiency on regeneration could be mitigated by the cytoprotective effects of necroptosis inhibition. Not to be forgotten that the regenerative process remains a consequence of previous degenerative events. Hence, the less degeneration, the less resulting regeneration. Therefore, any protection against myonecrosis is expected to hide any putative default on regeneration.

In mdx mice, myofibre necrosis does not lead to a dramatic lack of function of the limb, respiratory or cardiac muscles^29^. On another hand, pathology is dramatically exacerbated in R-DMDdel52 rats with a reduced life expectancy and cardiomyopathy functionally assessed in young adults^19^. Phospho-Ser161 RIPK1 is strongly upregulated in nuclei from cardiomyocytes, and the RIPK1/3 axis is largely overexpressed, suggesting ongoing necroptosis activation. In mdx mice, the heart pathogenic phenotype becomes detectable using M-mode echocardiography only after 10-12 months of age compared to wild-type mice. Heart weight, histological features, and analysis can discriminate healthy from dystrophic aged hearts^25,30,31^. Necroptosis depletion reduced heart weight, and biomarkers of hypertrophic cardiomyopathy such as *Bnp* and *Myh7* transcripts^32,33^ and fibrosis (Figures 4). BNP is upregulated in mdx compared to C57BL/10 control mice and restored to baseline in mdx mice treated by rAAV9-mediated microdystrophin^34^. The RIPK1/3 axis may therefore represent an interesting therapeutic target to counteract the development of cardiomyopathy in DMD. The role of canonical necroptosis has been found in cardiovascular diseases, such as atherosclerosis, ischemia-reperfusion injuries, myocardial infarction, myocarditis and chemically-induced cardiomyopathy^35^. We failed to observe a significant decrease in cardiomyocyte demise and necroinflammation. This is likely due to rare cell death events affecting the cardiomyocytes of mdx mice at this age. Indeed, only a minority of mdx hearts showed ongoing active necroinflammation area (Figure 4f). However, a reduction of cardiac hypertrophy linked with a reduction of fibrosis deposition suggests, over time, less accumulation of myocardial cell death susceptible to be replaced by collagen and overall improved tissue homeostasis.

In conclusion, using dogs, rats and mice models of DMD pathogenesis, we found that RIPK3 participates in the pathogenesis of mdx mice at different levels, mediating necrosis and fibrosis accumulation in locomotor, respiratory, and cardiac muscles without evidence of a detrimental effect on skeletal muscle regeneration. This study, therefore, suggests that necroptosis prevention can be beneficial both to skeletal muscle and heart phenotype in DMD conditions and paves the way for considering necroptosis as a promising therapeutic target for DMD.

## Acknowledgements

We thank Dr V. M. Dixit for *Ripk3* knockout mice. We also thank J. Ternacle, B. Drayton Libotte, L. Guillaud and X. Decrouy for technical assistance and C. Laisne and D. Gelperowic for taking good care of animals, and the VOUSH group for fruitful discussions. This work was supported by the Association Française contre les Myopathies (AFM-Téléthon) through the Translamuscle I (#19507) and Translamuscle II (#22946) programs.

## Supplementary data

**Supplementary table 1:** M-Mode echocardiographic values.

|  | Mouse# | Measured values |  |  |  |  |  |  |  |  |  | Calculated values |  |  |  |
| --- | --- | --- | --- | --- | --- | --- | --- | --- | --- | --- | --- | --- | --- | --- | --- |
|  |  | Tibial length (mm) | Normalizing factor | HR, beats/min | IVSd, mm | IVSs, mm | LVIDd, mm | LVIDs, mm | PWd, mm | PWs, mm | SRant, s-1 | LV RWT, mm | LV mass, mg | LVFS, % | LVEF, % |
| mdx | 1 | 21.1 | 0.912 | 523 | 0.78 | 1.45 | 3.81 | 1.85 | 0.74 | 1.68 | 26 | 0.40 | 95.75 | 51 | 88 |
|  | 2 | 18.2 | 1.056 | 605 | 0.94 | 1.64 | 3.92 | 1.70 | 0.76 | 1.49 | 29 | 0.43 | 117.26 | 57 | 91 |
|  | 3 | 19.2 | 1.006 | 513 | 0.85 | 1.60 | 3.60 | 1.56 | 0.98 | 1.48 | 27 | 0.51 | 113.44 | 57 | 91 |
|  | 4 | 19.1 | 1.011 | 535 | 0.86 | 1.88 | 3.52 | 1.63 | 1.13 | 1.88 | 25 | 0.57 | 123.85 | 54 | 89 |
|  | 5 | 18.8 | 1.027 | 605 | 0.85 | 1.13 | 3.72 | 2.38 | 0.85 | 1.31 | 29 | 0.46 | 108.05 | 36 | 72 |
|  | Mean | 19.3 | 1.0 | 556.2 | 0.9 | 1.5 | 3.7 | 1.8 | 0.9 | 1.6 | 27.2 | 0.5 | 111.7 | 50.9 | 86.2 |
|  | SD | 1.10 | 0.05 | 45.22 | 0.06 | 0.28 | 0.16 | 0.33 | 0.16 | 0.22 | 1.79 | 0.07 | 10.60 | 8.65 | 8.04 |
| mdxRipk3 -/- | 1 | 18.1 | 0.932 | 545 | 0.74 | 1.43 | 3.28 | 1.48 | 0.75 | 1.26 | 24 | 0.45 | 101.18 | 55 | 91 |
|  | 2 | 17.6 | 0.961 | 503 | 0.89 | 1.75 | 3.16 | 1.11 | 0.78 | 1.53 | 26 | 0.53 | 112.28 | 65 | 95 |
|  | 3 | 17.8 | 0.950 | 609 | 0.79 | 1.56 | 2.76 | 1.03 | 0.85 | 1.30 | 24 | 0.59 | 87.89 | 63 | 94 |
|  | 4 | 20.7 | 0.816 | 615 | 0.59 | 1.17 | 2.41 | 1.34 | 0.69 | 1.08 | 27 | 0.53 | 50.19 | 45 | 82 |
|  | 5 | 19.7 | 0.855 | 650 | 0.74 | 1.17 | 2.21 | 1.19 | 0.76 | 1.06 | 27 | 0.68 | 55.91 | 46 | 84 |
|  | 6 | 19.2 | 0.879 | 640 | 0.76 | 1.22 | 2.60 | 1.46 | 0.78 | 1.17 | 23 | 0.59 | 74.12 | 44 | 82 |
|  | 7 | 18.4 | 0.915 | 634 | 0.76 | 1.35 | 2.56 | 1.46 | 0.76 | 1.38 | 25 | 0.59 | 70.62 | 43 | 80 |
|  | Mean | 18.8 | 0.9 | 599.4 | 0.8 | 1.4 | 2.7 | 1.3 | 0.8 | 1.3 | 25.1 | 0.6 | 78.9 | 51.5 | 86.9 |
|  | SD | 1.15 | 0.05 | 54.76 | 0.09 | 0.22 | 0.39 | 0.18 | 0.05 | 0.17 | 1.57 | 0.07 | 22.86 | 9.25 | 6.28 |

